# Integrating metabolic scaling and coexistence theories

**DOI:** 10.1101/2023.02.28.530509

**Authors:** Serguei Saavedra, José Ignacio Arroyo, Jie Deng, Pablo A. Marquet, Christopher P. Kempes

## Abstract

Metabolic scaling theory has been pivotal in formalizing the expected energy expenditures across populations as a function of body size. Coexistence theory has provided a mathematization of the environmental conditions compatible with multispecies coexistence. Yet, it has been challenging to explain how observed community-wide patterns, such as the inverse relationship between population abundance density and body size, can be unified under both theories. Here, we provide the foundation for a tractable, scalable, and extendable framework to study the coexistence of resource-mediated competing populations as a function of their body size. For a given thermal domain and response, this integration reveals that the metabolically predicted 1/4 power dependence of carrying capacity of biomass density on body size can be understood as the average distribution of carrying capacities across feasible environmental conditions, especially for large communities. In line with empirical observations, our integration predicts that such average distribution leads to communities in which population biomass densities at equilibrium are independent from body size, and consequently, population abundance densities are inversely related to body size. This integration opens new opportunities to increase our understanding of how metabolic scaling relationships at the population level can shape processes at the community level under changing environments.

## Metabolic scaling theory

Metabolism is the sum of the total biochemical reactions governing the flow of energy in an individual (Kleiber, 1932). Therefore, knowing how and why metabolism changes across individuals has become central in our understanding of the energy requirements driving the sustainability of different forms of life on Earth (Arroyo et al., 2022, Kempes et al., 2011, Maurer, 1996, Shestopaloff, 2024). In this line, metabolic scaling theory (Brown et al., 2004) has been instrumental in investigating whether the rate of energy expenditure in an individual scales with its body size (body mass) and temperature. The size and temperature dependence is expected as the result of the fractal-like design of the resource supply system and the kinetics of the metabolic reactions (Gillooly et al., 2001, West et al., 1997). These dependencies are described by simple equations. On the one hand, the size dependence is typically allometric (not proportional) power-law relationships of the form *y* = *a* · *x*^*β*^, where *a* is the constant of proportionality (normalization constant), and the sign and magnitude of the exponent (power) *β* phenomenologically represent the physico-chemical constraints operating at the individual level (Gillooly et al., 2001, 2002, West et al., 1997). On the other hand, the temperature response (*f* (*T*)) can be either exponential or unimodal (Arroyo et al., 2022, Dell et al., 2011, Gillooly et al., 2001, 2002, West et al., 1997).

Put together, it has been shown (Gillooly et al., 2001, 2002, West et al., 1997) that the metabolic rate (*b*_*i*_) of an individual *i* is expected to scale with its body size (*m*_*i*_) under a given temperature (*T*) as

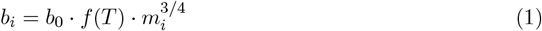

(dimensions [mass] · [time]^−1^), where *b*_0_ is a taxon-specific normalization constant (dimensions [mass]^1*/*4^[time]^−1^), and *f* (*T*) is a generic thermal response (dimensionless). Eq. 1 reveals that larger individuals are expected to be characterized by higher energy requirements per unit of time. These requirements, however, are expected to decrease non-linearly per unit of mass as 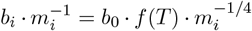 (known as mass-specific metabolism, dimension [time]^−1^), revealing the economies of body plans (Arroyo et al., 2022, Savage et al., 2004).

In community ecology (Vellend, 2016), populations (rather than individuals) are typically the basic units of living matter organization (Odum and Barrett, 2005, Vellend, 2016). Assuming a fixed growth rate of a population *i* under a constant environment, and that a stable age and size distribution have been reached, the average value of mass per individual (*M*_*i*_) becomes time independent and the average metabolic rate (Eq. 1) of population *i* can be written as 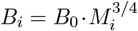 (dimensions [mass]·[time]^−1^) (Savage et al., 2004), where *B*_0_ is a now an effective parameter for a given temperature and thermal response (dimension [mass]^1*/*4^ · [time]^−1^). These metabolic scaling relationships have been confirmed by multiple empirical data across different taxa (Belgrano et al., 2002, Bernhardt et al., 2018, Damuth, 1987). However, the specific parameter values characterizing the metabolic dependence on temperature and size can vary as a function of the species and taxonomic group, respectively (Arroyo et al., 2022, Dell et al., 2011, Gillooly et al., 2001). For instance, it has been shown that for metazoans, protists, and prokaryotes, the scaling dependence of metabolic rate on body size ranges from sub-linear, linear, to super-linear, respectively (DeLong et al., 2010).

Similarly, it can then be predicted (Gillooly et al., 2002, Savage et al., 2004) that the maximum generation time (or lifespan) of an individual from population *i* becomes proportional to the scaling relationship 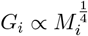 (dimension [time]). Thus, the reciprocal of maximum generation time (i.e., mortality rate) is expected to be proportionally related to the maximum growth rate of an individual in population *i* and can be written as

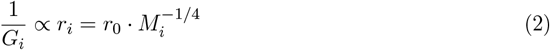

(dimension [time]^−1^), where *r*_0_ is an effective parameter for a given temperature and thermal response (dimension [mass]^1*/*4^ · [time]^−1^). These relationships have been used to show how the metabolic processes at the level of individuals can affect the metabolic and ecological processes at the level of populations (Savage et al., 2004).

Moreover, assuming that individuals from a population with different body sizes capture, on average, the same amount of resources or energy in their environment (i.e., the energetic equivalence rule (Damuth, 1987)), it has been shown (Savage et al., 2004) that it is possible to derive the mass and temperature dependence of a population’s carrying capacity of biomass density. This is the predicted, maximum, biomass density of a population in isolation that can be sustained and can be written as

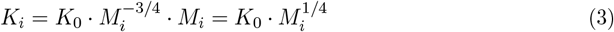

(dimensions [mass]·[area]^−1^), where *K*_0_ is an effective parameter (dimensions [mass]^3*/*4^·[area]^−1^). That is, under a given temperature and thermal response, the maximum biomass of a population in isolation per unit area is expected to increase with size. Note that this is different from the carrying capacity of abundance density 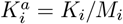 (dimensions [individuals]·[area]^−1^), which is expected to decrease with body size as 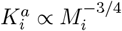 (Damuth, 1987).

Understanding how these different scaling relationships at the individual (or population level) are linked to ecological processes at the community level (the collection of different interacting populations in a given place and time) is challenging, and while there has been significant empirical and theoretical work using metabolic scaling theory at the community level (Basset and Angelis, 2007, Brose et al., 2006, Marquet et al., 2004, 1990, 2005, Tabi et al., 2023, Vasseur and McCann, 2005, Yeakel et al., 2018, Yodzis and Innes, 1992), several important challenges for theoretical integration and simplification still remain open. A clear example is the relationship between the distribution of body sizes and the coexistence of multiple populations, which has been typically studied for a specific set of model parameters (Brose et al., 2006, Yodzis and Innes, 1992). As a consequence, it is unclear the extent to which both the coexistence of multiple populations as well as their biomass densities depend on the distribution of body sizes (Bideault et al., 2021, Brose et al., 2006, DeLong et al., 2010, Hatton et al., 2019, Savage et al., 2004). Similarly, there is still theoretical work to be done in order to increase our understanding of how metabolic scaling relationships at the individual or population levels affect coexistence processes at the community level (Arim et al., 2007, Basset and Angelis, 2007, Hatton et al., 2024, Savage et al., 2004).

## Coexistence theory

In general, coexistence theory in ecology (Chesson, 2000, MacArthur and Levins, 1967, Saavedra et al., 2017, Vandermeer, 1970) aims to investigate the possibility that a community 𝒮(**A**), formed by |𝒮| populations and characterized by an interaction matrix (**A**), can persist under different environmental contexts. Although not exclusively, this analysis has been mainly performed under generalized Lotka-Volterra dynamics (Case, 2000), which in line with metabolic scaling theory (Brown et al., 2004), can be derived from thermodynamics principles, from principles of conservation of mass and energy, and from chemical kinetics in large populations (Logofet, 1993, Lotka, 1920, Michaelian, 2005, Täuber, 2011). In particular, in the Lotka-Volterra competition model (Arditi et al., 2021, Medeiros et al., 2021a), the per capita growth rate of a population’s biomass density can be written as

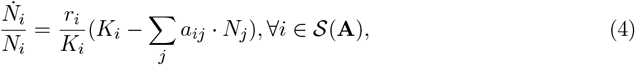

(dimension [time]^−1^), where *N*_*i*_ is the population biomass density (dimensions [mass]·[area]^−1^) and 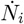 is the time derivative (dimensions [mass]·[area]^−1^·[time]^−1^). Here, *r*_*i*_ *>* 0 is the maximum intrinsic growth rate (dimension [time]^−1^), *K*_*i*_ *>* 0 is the carrying capacity of biomass density (dimensions [mass]·[area]^−1^). Then, **A** = (*a*_*ij*_) ≥ 0 ∈ ℝ^|𝒮|×|𝒮|^ is the per capita, time-independent, area-independent, competitive effect (dimensions [mass_*i*_]·[mass_*j*_]^−1^) of a population *j* on an individual of population *i*. Note that

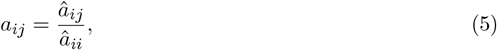

where *â*_*ij*_ and *â*_*ii*_ are the competitive effects across unit area (dimensions [time]^−1^·[mass_*j*_]^−1^·[area]) and self-regulation (dimensions [time]^−1^·[mass_*i*_]^−1^[area]), respectively. It follows that *a*_*ii*_ = 1 per definition. In other words, *a*_*ij*_ specifies the time-independent, area-independent rate of biomass converted from a population *i* to a population *j* (or resources used by *j* instead of *i*) (Logofet, 1993). Then, the carrying capacity of biomass density of a population *i* under a given environment can be expressed as a function of its intrinsic growth rate and self-regulation as

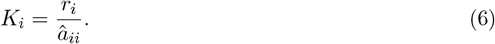

The necessary condition for persistence under Lotka-Volterra equilibrium dynamics (i.e., 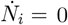) becomes the existence of positive solutions in population biomass densities (Hofbauer and Sigmund, 1998). This necessary condition is known as *feasibility* and takes the form of a linear equation in matrix notation (Saavedra, 2024, Vandermeer, 1970))

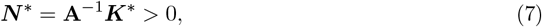

where the vector ***N*** ^*^ represents the distribution of biomass densities at equilibrium, **A**^−1^ corresponds to the inverse of the interaction matrix, and the vector ***K***^*^ is the distribution of carrying capacities of biomass densities compatible with the feasibility of community 𝒮(**A**). Notice that it is possible to multiply the vector ***K***^*^ by any positive scalar (*λ >* 0) and the feasibility condition (i.e., ***N*** ^*^ = **A**^−1^***K***^*^*λ >* 0) does not change. This is true because the relationship among carrying capacities of biomass densities within the community does not change (i.e., all values are either increased or decreased proportionally). Of course, if *λ* approaches zero, the loss of feasibility can happen due to stochasticity (Hofbauer and Sigmund, 1998). This implies that given an interaction matrix, the feasibility of a community is determined by the direction rather than the magnitude of the vector (Saavedra et al., 2017). In other words, feasibility depends on the relationship (or distribution) among carrying capacities of biomass densities within the community.

Importantly, there can be more than one compatible vector ***K***^*^. This possible set of vectors is known as the feasibility domain (*D*_*F*_) (Logofet, 1993, Saavedra et al., 2017). Formally, rearranging Eq. 7 and solving for ***K***^*^, this feasibility domain can be expressed as

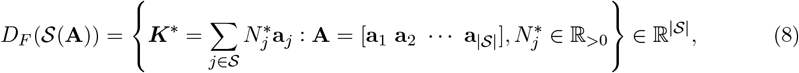

where **a**_*j*_ is the *j*^th^ column vector of the interaction matrix **A** (Logofet, 1993, Song and Saavedra, 2018). Each column *j* represents the time-independent, area-independent, competitive effect of population *j* on all other populations (including itself). The feasibility domain specifies all the possible distributions (vector directions) of *K* (Eq. 6) that are compatible with the interaction matrix **A** as written in Eq. (7). Geometrically, the feasibility domain is the convex hull of the |𝒮| spanning vectors in |𝒮|-dimensional space (Fig. 1) (Deng et al., 2023, 2022, Song et al., 2018*b*). Overall, the interaction matrix **A** determines the range of the feasibility domain. The stronger the mean competition in a community (⟨*a*_*ij*_⟩), the smaller the range of the feasibility domain (Saavedra et al., 2017).

**Figure 1.**
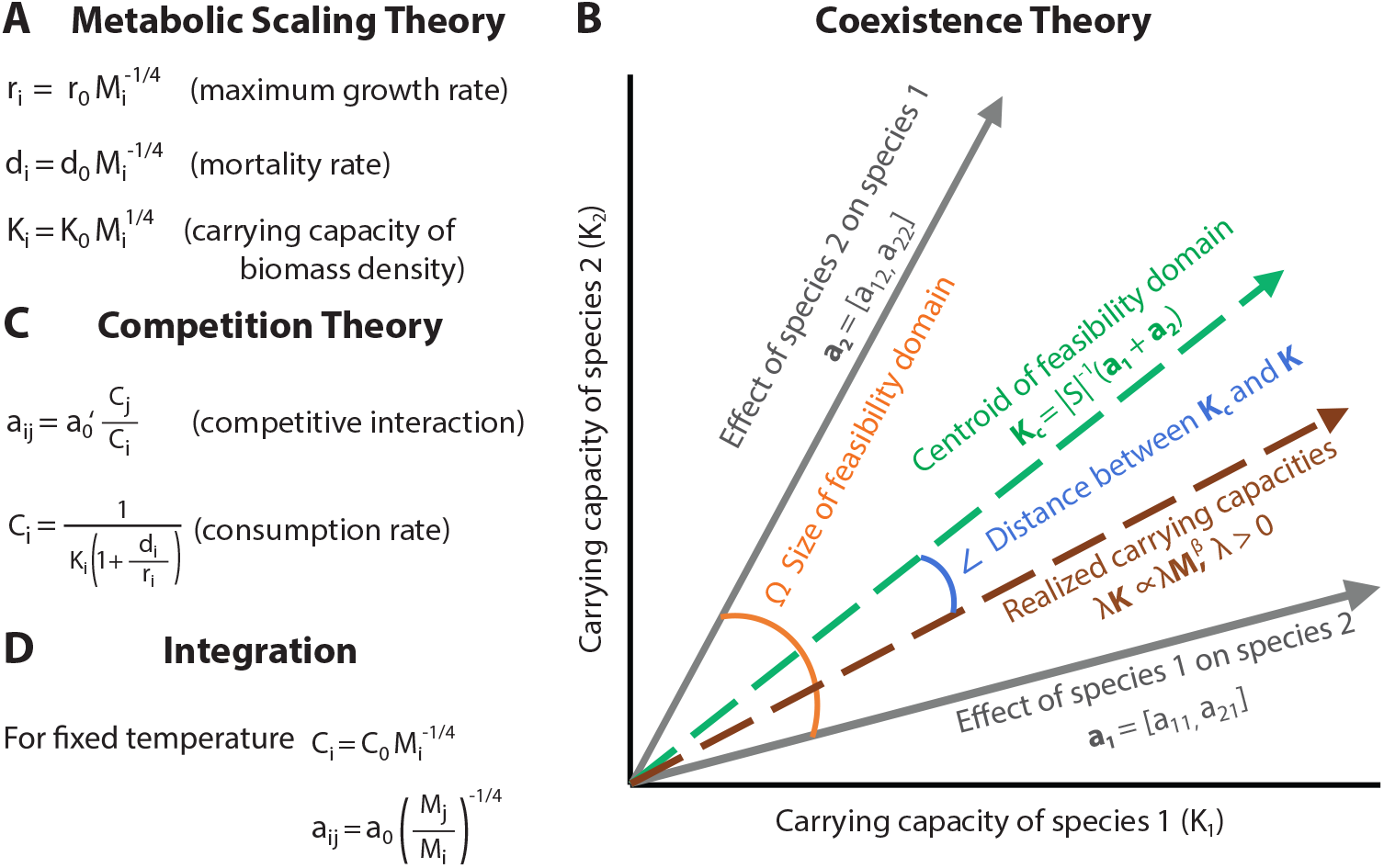
A graphical integration of metabolic scaling and coexistence theories. (**A**) Following metabolic scaling theory, for a given temperature and thermal response, maximum growth rate, mortality rate, and carrying capacities of biomass densities are predicted to scale as a function of body mass. (**B**) Following coexistence theory and under Lotka-Volterra competition dynamics (see main text), the feasibility (i.e., positive biomass densities at equilibrium) of a community will be defined by having the relationship (vector direction) among carrying capacities inside the domain constrained by species interactions. As an illustrative example, we show the parameter space (Cartesian coordinate system) of carrying capacities of biomass densities *K*_*i*_ of a community with |*S*| = 2 populations. The outer solid gray arrows correspond to the column vectors of the interaction matrix (**A** = [**a**_**1**_; **a**_**2**_]). The angle (orange large curve) formed by the two column vectors corresponds to the feasibility domain *D*_*F*_ (𝒮 (**A**)) (Eq. 8). Inside the two column vectors, any direction of carrying capacities **K** is compatible with positive biomass densities at equilibrium (Eq. 7). In this sense, the direction (rather than the magnitude) defines the feasibility of a community. The centered green vector depicts the geometric centroid of the feasibility domain (Eq. 9, **K**_**c**_), representing the average distribution of carrying capacities of biomass densities across feasible environmental conditions for large communities. While the elements of ***K***_***c***_(**A**) can be different from each other, the centroid corresponds to the carrying capacities that provide the solution where all biomass densities are equal. (**C**) Following competition theory (see main text), we can define the time-independent, area-independent competitive effect (*a*_*ij*_) of species *j* on *i* as a function of the average consumption rates, which can also be expressed as a function the maximum growth rate, mortality rate, and carrying capacity of biomass densities. (**D**) Integrating theories (see main text), for a fixed temperature, we can then express both the average consumption rates and the competitive effect as a power law function of body size.

For a given community characterized by a particular interaction matrix, the location of a given vector ***K***^*^ inside the feasibility domain provides information about a given set of environmental conditions compatible with the feasibility of such community (i.e., ***N*** ^*^ *>* 0). Formally, the geometric centroid of the feasibility domain (Fig. 1) can be understood as the average distribution of carrying capacities of biomass densities across all feasible environments. This statement makes the assumption that the elements of the vector of carrying capacities of biomass densities are independent from each other and free to vary due to external factors other than temperature, such as long-term changes in species habitats. Formally, the geometric centroid can be calculated as (Medeiros et al., 2021b, Song and Saavedra, 2018, Tabi et al., 2020)

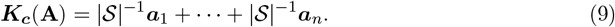

Moreover, by mathematical definition, the centroid ***K***_***c***_(**A**) yields the solution (Rohr et al., 2016)

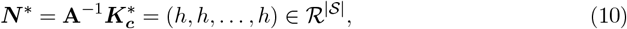

where *h >* 0 is a positive constant. That is, while the elements of the vector of carrying capacities of biomass densities defining the centroid of the feasibility domain can be different from each other, the populations’ biomass densities become the same at equilibrium and independent from body size.

## Integrating theories

The link between metabolic scaling and population dynamics has been performed under different frameworks, each answering specific ecological questions (Basset and Angelis, 2007, Brose et al., 2006, Campillay-Llanos et al., 2022, Harte et al., 2008, Parain et al., 2019, Vasseur and McCann, 2005, Yeakel et al., 2018, Yodzis and Innes, 1992, Zaoli et al., 2017). Here, we provide a tractable, scalable, and extendable framework to study the effect of body size on the feasibility of resource-mediated competing populations. Specifically, we aim to investigate the possibility that a community of competing populations characterized by a given distribution of body sizes can coexist under different environmental contexts. To this end, our framework is based on Lotka-Volterra competition dynamics (Vandermeer and Goldberg, 2013).

Under resource-mediated competition models, it is assumed that the dynamics of resources are faster than that of consumers. While this is a strong assumption, classic and recent work has shown promising approximations to empirical competition dynamics (MacArthur and Levins, 1967, Medeiros et al., 2021*a*, Parain et al., 2019, Song et al., 2018*a*). Following this premise, time-independent, area-independent, competitive interactions (*a*_*ij*_) can be written as a function of the average consumption rates of consumers (Fig. 1, see Appendix S1 for details):

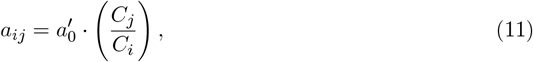

where *C*_*i*_ corresponds to the time-independent, average consumption rate for a population *i* across unit area (dimensions [mass_*i*_]^−1^·[area]), and *a*′_0_ is an *effective* parameter representing the overall effect of intrinsic properties (dimensionless), such as consumers’ conversion rates (Parain et al., 2019). Similarly, following single population models (Deng et al., 2024), the time-independent, average consumption rate of consumer *i* can be expressed as a function of its demographics (Fig. 1, see Appendix S1 for details):

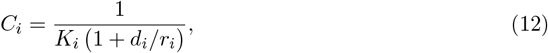

where *d*_*i*_ denotes the mortality rate (dimension [time]^−1^), and *K*_*i*_ and *r*_*i*_ represent again the carrying capacities of biomass densities and maximum growth rate of consumer *i*, respectively.

Then, we can integrate metabolic scaling theory (Eqs. 2-3) and coexistence theory (Eq. 12) to write

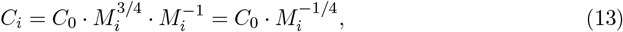

where *C*_0_ is a new effective parameter (dimensions [mass]^−3*/*4^·[area]). Note that studies have also found different scaling dependencies between consumption rate and body size (Pawar et al., 2012). As we will show below, these differences do not affect our conclusions. Thus, for a fixed temperature and thermal response, we can integrate Eqs. 11 and 13 as

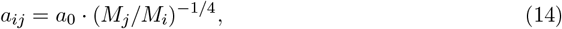

where *a*_0_ is yet again a new effective parameter ([mass_*i*_]^3*/*4^·[mass_*j*_]^−3*/*4^). In sum, for a given thermal domain and response, it is expected that the higher the average body size of population *j* relative to that of population *i*, the lower the time-independent, area-independent, competitive effect.

Next, we investigated the link between the feasibility of a community and the distribution of body sizes for a fixed temperature in metabolically-generated competition matrices **A**. In particular, we systematically studied the scaling relationship *β* between the vector of carrying capacities of biomass densities and the vector of body sizes (***K*** = ***K***_**0**_ · ***M*** ^*β*^) that makes ***K*** close to the centroid of the feasibility domain (***K***_***c***_(**A**)). Recall that the centroid represents the average of carrying capacities of biomass densities across feasible environmental conditions, as expressed in Eq. 7.

Specifically, following Eq. 14, we generated an ensemble of 10^4^ competition matrices **A** characterizing the time-independent, area-independent, competitive effects (*a*_*ij*_) between 50 populations (larger dimensions yield similar conclusions, see Appendix S1). Each matrix was formed by drawing *M*_*i*_ values independently from a lognormal distribution *LN* (0, 2). These distributions can change without affecting the qualitative results. Following previous work (Bunin, 2017, Dougoud et al., 2018), we set the effective parameter of competitive effects to *a*_0_ = |𝒮|^−1*/*2^ for *i* ≠ *j*, otherwise *a*_0_ = 1. This assumption follows the rationale that empirical interactions tend to be weak, stabilizing communities (Gellner et al., 2023, McCann et al., 1998). The greater the value of *a*_0_, the greater the overall competition, and the smaller the range of the feasibility domain. Then, for regular intervals between *β* ∈ [−2, 2] (used as representative values), we calculated the average distance between ***K***_***c***_(**A**) and ***K*** as (Medeiros et al., 2021*b*, Saavedra et al., 2017)

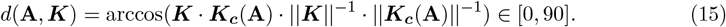

Note that the normalization constant *K*_0_ of carrying capacities of biomass densities does not affect this distance and can be omitted from the equation (Rohr et al., 2016).

Figure 2 reveals that the vector of carrying capacities of biomass densities described by the scaling relationship 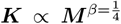 gets close to the centroid of the feasibility domain of the metabolically-generated competition matrices for a given thermal domain and response. For few competing species (e.g., two), it will be exactly the centroid only for a particular value of *a*_0_ (see Appendix S1). This vector tends to get asymptotically closer to the centroid of the feasibility domain the larger the community (see Appendix S1 for numerical analysis). This reveals that the expected 1/4 power dependence of carrying capacities of biomass densities on body size can be understood as the average distribution of carrying capacities across feasible environmental conditions, especially for large communities. Furthermore, because the centroid represents the condition under which all populations have the same biomass densities at equilibrium (Eq. 10), dividing these biomass densities (***N*** ^*\**^) by the corresponding average body size ***M***, the number of individuals per area (***I***) become inversely related to body size (***I*** ∝ ***M*** ^−1^). This mathematical fact is independent from the scaling exponent defining the dependency of consumption rate and body size (Pawar et al., 2012). This implies that our framework is in agreement with empirical observations showing that, at the community level (or over a much larger size range of life), a population’s abundance density scales inversely with body size (Hatton et al., 2019, Marquet et al., 1990, Segura and Perera, 2023).

**Figure 2.**
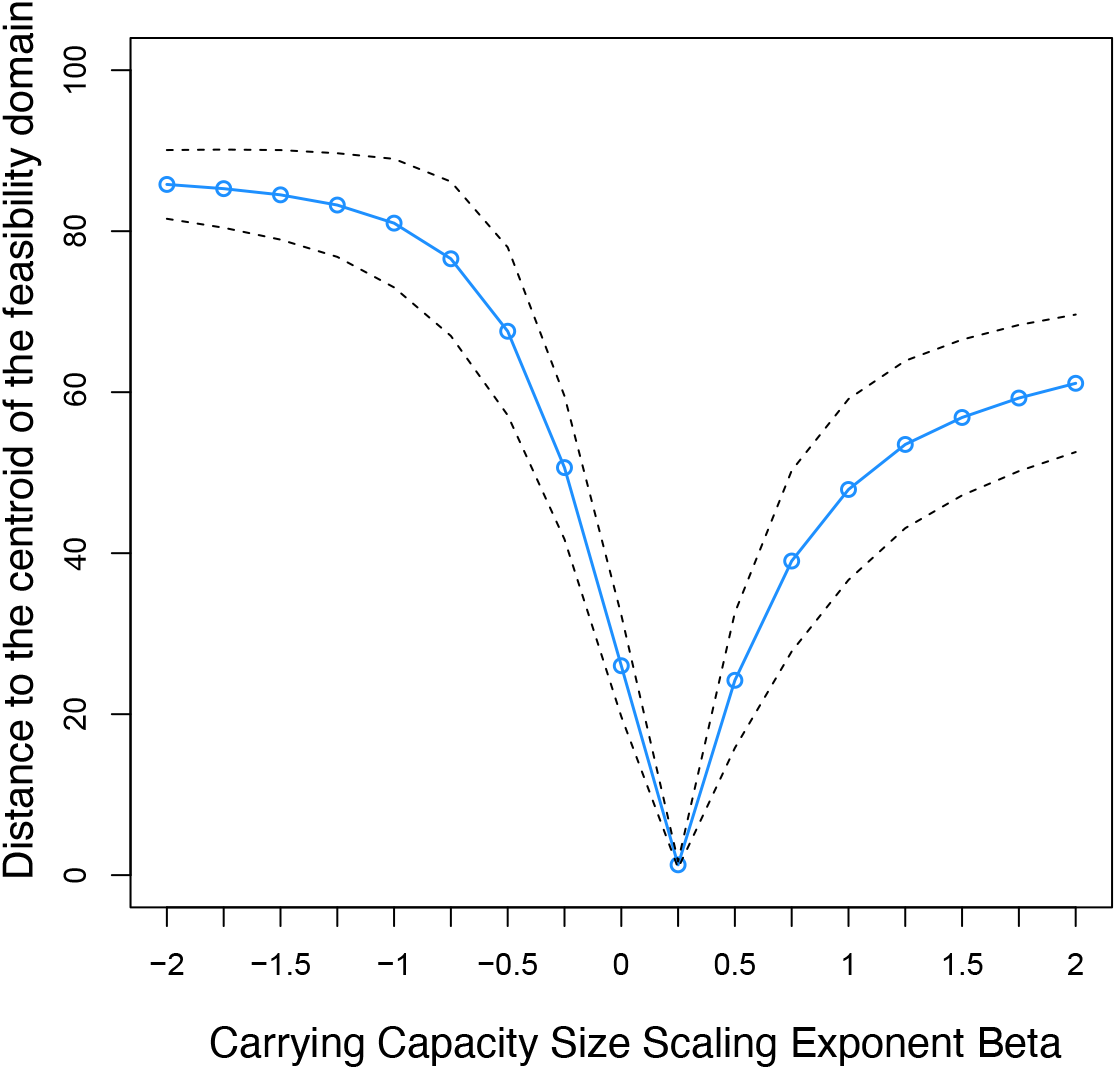
Theoretical predictions. For an ensemble of 10^6^ communities with |*S* = 50v| populations, the blue solid line shows the average distance (Eq. 27, blue small curve in Figure 1) between the centroid of the feasibility domain **K**_**c**_ and the possible vectors of carrying capacities of biomass densities (***K***∝***M*** ^*β*^) as a function of the power exponent beta (*β*∈ [−2, 2]) describing the dependence of carrying capacities on body size. The black dashed lines correspond to the average plus minus two standard deviations. For each community, the interaction matrix (Eq. 11) is defined by *α* = −1*/*4 and *a*_0_ = | *S*| ^−1*/*2^. The values of *M*_*i*_ are drawn independently from a lognormal distribution *LN* (0, 2). For large communities, *β* = 1*/*4 tends to be the closest to the centroid (see Supplementary Information for more details).

## Concluding remarks

We have proposed an tractable, scalable, and extendable integration of metabolic scaling theory (Brown et al., 2004) and coexistence theories (Chesson, 2000, MacArthur and Levins, 1967, Saavedra et al., 2017, Vandermeer, 1970) in order to systematically link metabolic scaling relationships at the population level to community coexistence (Deng et al., 2021, 2022, Saavedra et al., 2020). Our framework has been based on understanding how body-size constraints, operating on competitive effects, modulate the parameter space of carrying capacities of biomass densities compatible with multispecies coexistence. This integration has allowed us to show that the metabolically predicted 1/4 power dependence of carrying capacities of biomass densities on body size can be understood as the average distribution of biomass carrying capacities across feasible environmental conditions, especially for large communities. Additionally, in line with empirical observations (Hatton et al., 2019), our integration conforms to the expectation that biomass densities at equilibrium are independent from body size, and consequently, to abundance densities at equilibrium that are inversely related to body size. Therefore, our framework can be used to link competitive effects to metabolism and, consequently, to body size, helping to capture important theoretical derivations and empirical observations (Hatton et al., 2019, Savage et al., 2004).

It is worth mentioning that our efforts to integrate metabolic scaling and coexistence theories have been simplified under a formalism that uses a set of relevant assumptions. In particular, we have focused on resource-mediated competition assuming that the dynamics of resources are faster than that of consumers. While there is a strong tradition on this type of competition systems, this does not take into account other type of much richer dynamics, such as as trophic interactions. However, previous efforts on these forms of dynamics (Parain et al., 2019, Vasseur and McCann, 2005) can be compatible with our framework (Saavedra et al., 2016). Similarly, we have exclusively focused on feasibility (the necessary condition for species coexistence under equilibrium dynamics) without systematically investigating dynamical stability (Song and Saavedra, 2018). For instance, future work can expand our work in order to investigate the role of life-history traits acting on interspecific effects in shaping both the feasibility and dynamical stability of communities. Additionally, while our results on the distance to the centroid of the feasibility domain are valid for a fixed thermal domain and response, feasibility itself can change as a function of temperature via changes in interspecific effects. Thus, for cases where the interest is to predict how a population or community will respond to temperature, it would be useful to use a model that accounts for the curvature of temperature responses (Arroyo et al., 2022, Dell et al., 2011). This alternative model can be adapted to our integration (by separating the role of temperature from the effective parameters) to explore temperature-related questions, such as how the variation in optimum temperatures of populations in a community would affect its feasibility or how global warming may alter communities. We believe the proposed integration can open up new opportunities to increase our understanding of how metabolic scaling relationships at the individual and population levels shape processes at the community level.

## Acknowledgments

We would like to thank the Santa Fe Institute scaling and NETI working groups, Lucas Medeiros, Rudolf Rohr, Chuliang Song, Andrea Tabi, and Yuguang Yang for insightful comments that led to improving this work. SS acknowledges funding from MathWorks,

MIT Google Program for Computing Innovation, and the support by the National Science Foundation under Grant No. DEB-2436069. JD acknowledges funding from MathWorks and the Royal Society through the Newton International Fellowship. PAM acknowledges support from Centro de Modelamiento Matemático (CMM), Grant FB210005, BASAL funds for centers of excellence from ANID-Chile, and Projecto Exploración 13220168.

## Appendix S1

### Integrating metabolic scaling and coexistence theories

The estimation of time-independent, area-independent, competitive interactions *a*_*ij*_ is derived by translating classic consumer-resource dynamics into Lotka-Volterra competition dynamics following Vandermeer and Goldberg (2013).

Let us assume that there are two self-replicating resources:

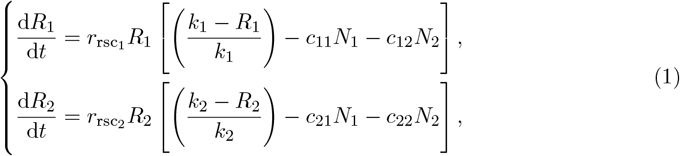

where *R*_*i*_ is the biomass density of the resource *i* (dimensions [mass]·[area]^−1^), 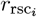 is the intrinsic rate of increase of resource *i* (dimension [time]^−1^), *k*_*i*_ is the carrying capacity of biomass density of the resource *i* (dimensions [mass]·[area]^−1^), *c*_*ij*_ is the time-independent rate at which the resource *i* is consumed by the *j*^th^ consumer across unit area (dimensions [mass_*j*_]^−1^·[area]), and *N*_*i*_ is the population biomass density of the *i*^th^ consumer (dimensions [mass]·[area]^−1^).

Then, considering two consumers, we can write the consumer equations:

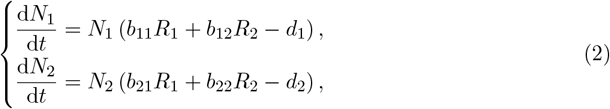

where *b*_*ij*_ is the relative conversion of a unit of resource *j* by consumer *i* across unit area per time (dimensions [mass_*j*_]^−1^ · [time]^−1^·[area]), and *d*_*i*_ is the death rate of the *i*^th^ consumer (dimension [time]^−1^). Note that *b*_*ij*_ is different from mass transformation efficiency.

Assuming that the dynamics of the resources occur significantly faster than those of the consumers (MacArthur and Levins, 1967), we can equate the derivatives of Eqs. (1) to zero and solve for the equilibrium values 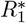 and 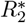 of the two resources:

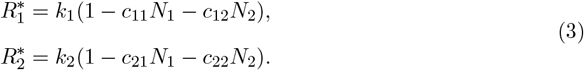

Then, we can substitute *R*_1_ and *R*_2_ in Eqs. (2) by 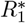 and 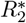, respectively. That is,

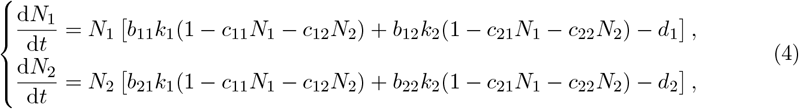

which can also be written as

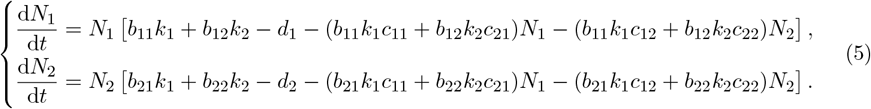

Multiplying and dividing the two equations in Eqs. (5) by

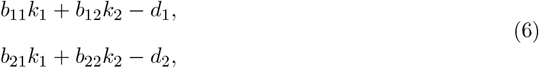

respectively, we obtain

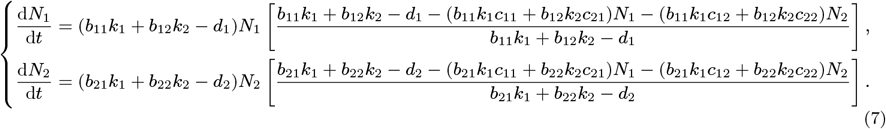

Again, dividing both the numerator and denominator of the fractions on the right sides of the two equations in Eqs. (7) by

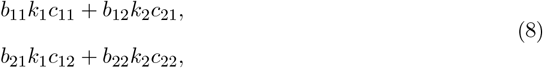

respectively, gives us

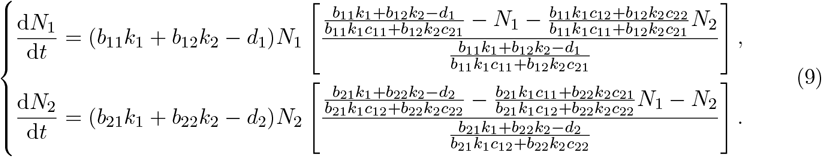

Essentially, Eqs. (9) are the Lotka-Volterra competition dynamics, which can be easily recognized if we make the following substitutions

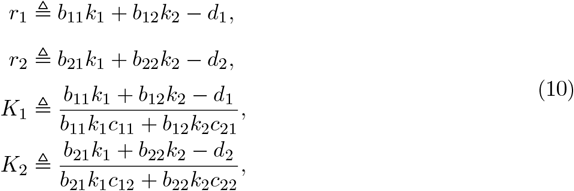

And

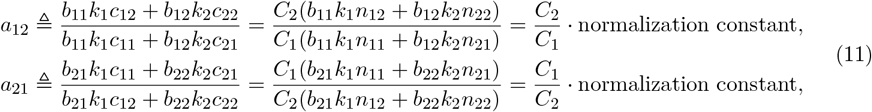

where *r*_*i*_ is the rate of natural increase in isolation of consumer *i* (dimension [time]^−1^), *K*_*i*_ is the carry capacity of consumer *i* biomass density in the absence of all competitors (dimensions [mass]·[time]^−1^), and *a*_*ij*_ is the time-independent, area-independent, competition coefficient measuring the competitive effect of consumer *j* on consumer *i* (dimensions [mass_*i*_]·[mass_*j*_]^−1^). Specifically, *C*_*j*_ is the time-independent average rate at which consumer *j* consumes resources across unit area (dimensions [mass]^−1^·[area]). Furthermore, it is typically observed that the relative conversion *b*_*ij*_ of a unit of resource *j* by consumer *i* and the carrying capacity *k*_*j*_ of resource *j* have limited variations for a given consumer *i* across different resources *j* and can be assumed constant for the sake of model simplification (Brown et al., 2004). Therefore, as indicated by Eq. (11), it is important to note that the competition coefficient *a*_*ij*_ is proportional to the ratio of consumption rates *C* between consumer *j* and consumer *i*. Formally, we can represent it as

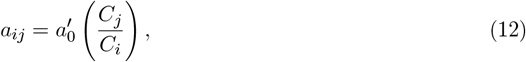

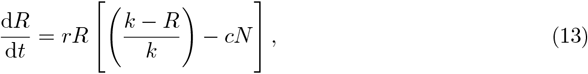

where *a*′_0_ is an *effective* parameter (dimensionless) representing the overall effect of intrinsic properties, such as consumers’ conversion rates (Parain et al., 2019).

To further estimate the time-independent average rate *C*_*i*_ at which consumer *i* consumes resources, we can simplify the previous scenario by considering a streamlined framework that involves only one consumer (also predator, either consumer *j* or consumer *i*) and one resource (also prey). Specifically, if the resource is self-replicating, then the resource equation with a logistic growth is where *R* is the biomass density of the resource (dimensions [mass]·[area]^−1^), *r* is its per capita rate of increase (dimension [time]^−1^), *k* is the carrying capacity of biomass density of the resource (dimensions [mass]·[area]^−1^), *c* is the time-independent rate at which the resource is consumed by the consumer across unit area (dimensions [area]·[mass]^−1^), and *N* is the biomass density of the consumer (dimensions [mass]·[area]^−1^).

Then, we can write the consumer equation as

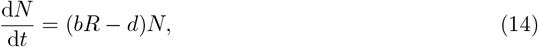

where *b* is the area-independent predation rate (dimensions [mass_*R*_]^−1^·[time]^−1^) and *d* is the death rate of the consumer (dimension [time]^−1^).

Assuming again that the resource dynamics are very fast compared to the consumer dynamics (MacArthur and Levins, 1967), we can solve the equilibrium value of R:

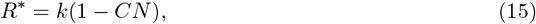

and then substitute *R*^*^ in place of *R* in the consumer equation Eq. (14). We obtain

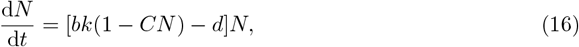

which can also be written as

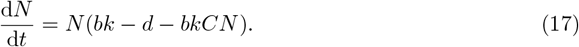

Multiplying and dividing Eq. (17) by (*bk* − *m*) gives us

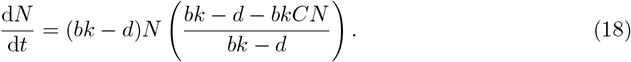

Next, dividing both the numerator and the denominator of the fraction on the right side of the Eq. (18) by *bkC*, we have

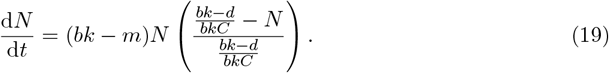

If we denote

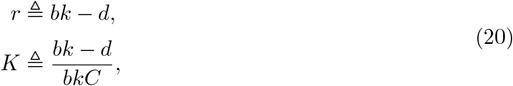

then Eq. (19) can be written as the classic population model with logistic growth:

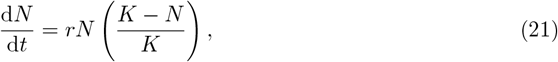

where again *r* is the rate of natural increase in isolation of the consumer (dimension [time]^−1^) and *K* is the carrying capacity of the consumer without competitors (dimensions [mass]·[area]^−1^).

Importantly, Eqs. (20) indicate that the time-independent consumption rate *C* increases monotonically with the intrinsic growth rate *r* (Deng et al., 2024). Formally, by substituting the expression for *bk* as (*r* + *d*) from the first equation into the second equation, we obtain

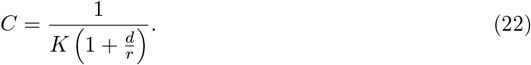

The time-independent consumption rate *C* (dimensions [area]·[mass]^−1^) exhibits a monotonically decreasing behavior with respect to the carrying capacity *K* (dimensions [mass]·[area]^−1^). Note that the relationship between carrying capacity *K* and intrinsic growth rate *r* appears to be complex and context-dependent (Marshall et al., 2023). Therefore, instead of establishing relationships between *r* and *K*, our primary focus lies on *K* due to its role in the feasibility of Lotka-Volterra competition dynamics. However, the focus can be shifted to *r* (Deng et al., 2024).

Following metabolic scaling theory (Eqs. 2-3 in main text), one can write the time-independent average consumption rate of population *i* (Eq. 35) as a function of body size (*M*_*i*_). Note that mortality rate (*d*_*i*_) is the reciprocal of generation time (*G*_*i*_) and proportional to growth rate (*r*_*i*_). For a fixed thermal domain and response, we can replace the parameters in Eq. 37 with the following equivalences: 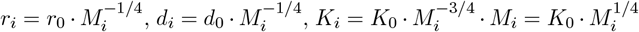,where *r*_0_, *d*_0_, and *K*_0_ are normalization constants. Thus, Eq. (35) can be expressed as:

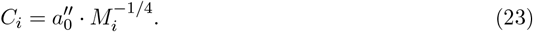

Eventually, by combining Eq. (12) with Eq. (36), we can derive an estimation of time-independent, area-independent, competitive interaction strength *a*_*ij*_ for a fixed thermal domain and response as:

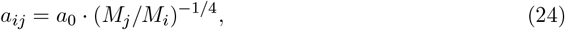

where 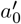, 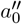 and *a*_0_ are normalization constants. Based on this estimation, we can generate the competition matrix **A** for the Lotka-Volterra competition dynamics in Eq. (4) in main text.

Under *a*_*ij*_ = *a*_0_·(*M*_*j*_*/M*_*i*_)^−1*/*4^ (Eq. 39), Figure 2 (main text) shows that if the carrying capacities of biomass densities are defined as 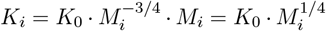,then they are the closest to the centroid of the feasibility domain. However, these carrying capacities ***K*** are expected to become exactly the centroid only under a specific value of *a*_0_. For example, following Eq. 39 and for two competing species, the maximum and minimum feasible carrying capacities for species 2 (corresponding to the spanning vectors of the feasibility domain in Fig. 1) can be defined as *K*_2max_ = (*a*_22_*/a*_12_)*K*_1_ and *K*_2min_ = (*a*_21_*/a*_11_)*K*_1_, where 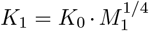 is the carrying capacity of biomass density of species 1 per metabolic scaling theory. Then, following Eq. 9, the carrying capacity for species 2 that belongs to the centroid of the feasibility domain becomes:

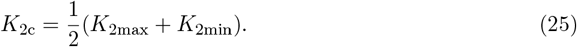

Note that in the centroid the distance beteween *K*_2*c*_ and *K*_1_ should be the absolute minimum, i.e., *K*_2*c*_*/K*_1_ = 1. Thus, diving Eq. 40 by the carrying capacities of biomass density of species 1 (*K*_1_) and rearranging it yields:

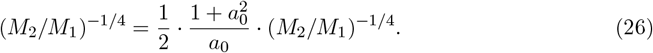

This implies that the equality is satisfied if and only if the factor 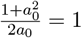.Since *a*_0_ also affects the dynamical stability of the community, future work can investigate the possible trade-offs between stability and feasibility under this framework. However, under numerical simulations, the larger the community, the closer this expression becomes the centroid of the feasibility domain.

The vector 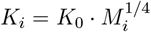 tends to get asymptotically closer to the centroid of the feasibility domain the larger the community (Fig S1). Specifically, following Eq. 14 in main text, we generated an ensemble of 10^4^ competition matrices **A** characterizing the time-independent, area-independent, competitive effects (*a*_*ij*_) between 10, 20, 50, 100, 200, 500, and 1000 populations. Each matrix was formed by drawing *M*_*i*_ values independently from a lognormal distribution *LN* (0, 2). These distributions can change without affecting the qualitative results. Following previous work (Bunin, 2017, Dougoud et al., 2018), we set the effective parameter of competitive effects to *a*_0_ = |𝒮|^−1*/*2^ for *i j*, otherwise *a*_0_ = 1. This assumption follows the rationale that empirical interactions tend to be weak, stabilizing communities (Gellner et al., 2023, McCann et al., 1998). The greater the value of *a*_0_, the greater the overall competition, and the smaller the range of the feasibility domain. Then, we calculated the average distance between ***K***_***c***_(**A**) and ***K*** as (Medeiros et al., 2021b, Saavedra et al., 2017)

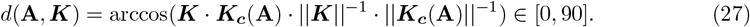

Note that the normalization constant *K*_0_ of carrying capacities of biomass densities does not affect this distance and can be omitted from the equation (Rohr et al., 2016). This numerical analysis reveal that the large the number of populations, the smaller the distance to the centroid.

**Figure 1.**
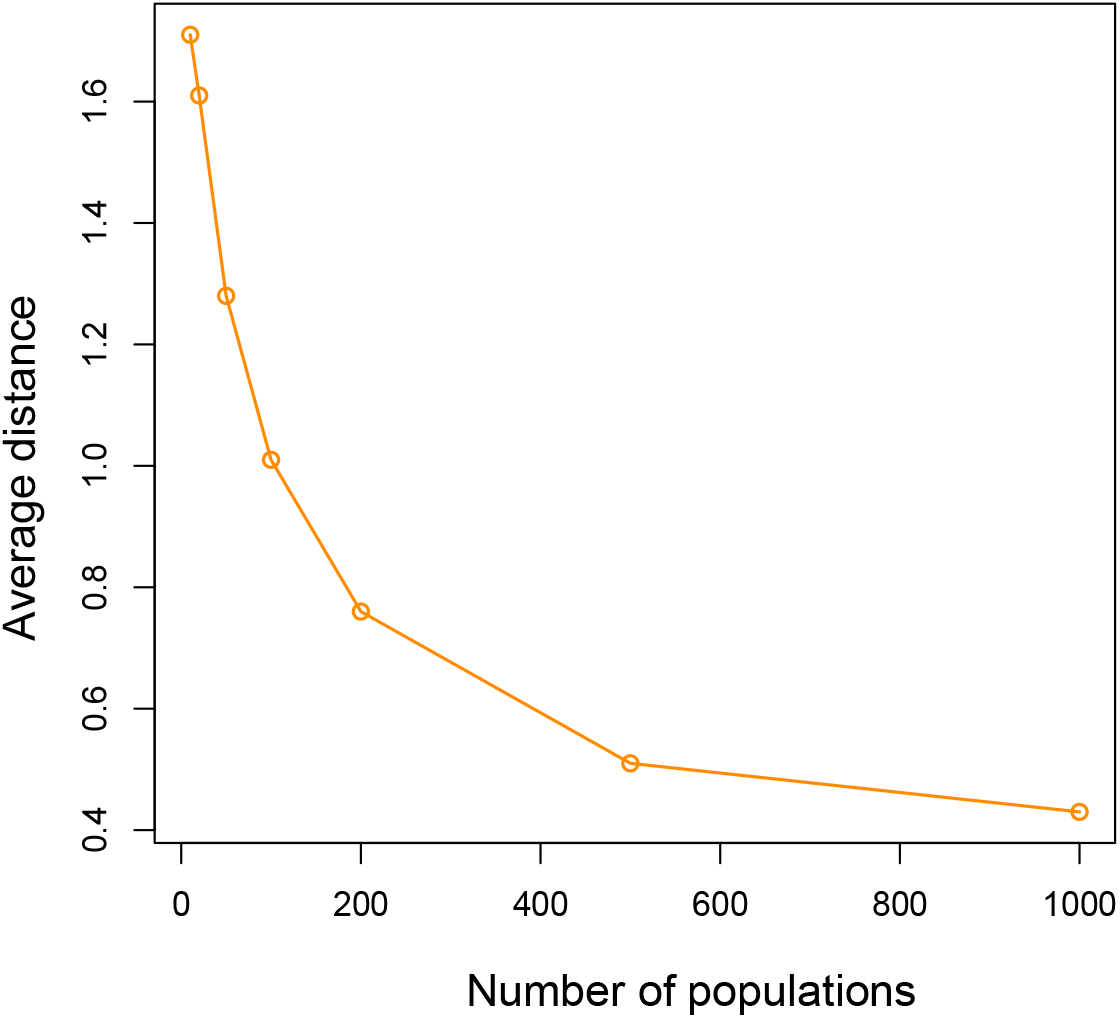
Distance to the centroid for large communities. For an ensemble of 10^4^ generated competition matrices with different number of populations, we show the average distance between the vector 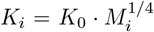 and the centroid of the feasibility domain. This numerical analysis reveal that the large the number of populations, the smaller the distance to the centroid.

